# Methods for distinguishing between protein-coding and long noncoding RNAs and the elusive biological purpose of translation of long noncoding RNAs

**DOI:** 10.1101/017889

**Authors:** Gali Housman, Igor Ulitsky

## Abstract

Long noncoding RNAs (lncRNAs) are a diverse class of RNAs with increasingly appreciated functions in vertebrates, yet much of their biology remains poorly understood. In particular, it is unclear to what extent the current catalog of over 10,000 distinct annotated lncRNAs is indeed devoid of genes coding for proteins. Here we review the available computational and experimental schemes for distinguishing between coding and noncoding transcripts and assess the conclusions from their recent genome-wide applications. We conclude that the model most consistent with available data is that a large number of mammalian lncRNAs undergo translation, but only a very small minority of such translation events result in stable and functional peptides. The outcome of the majority of the translation events and their potential biological purposes remain an intriguing topic for future investigation.

## Introduction – the gray area between coding and noncoding transcripts

Genome-wide surveys of transcription in mammalians [1-8] and more recently in other vertebrates [9-13] have shown that vertebrate genomes are pervasively transcribed and produce a large variety of processed transcripts, many of which do not overlap canonical genes. These include long noncoding RNAs (lncRNAs), which closely resemble mRNAs in that they possess an m^7^Gpppn cap at the 5′ end, a poly(A) tail at the 3′ end, and in most-cases undergo splicing [14]. The expression of many lncRNAs has been shown to be altered in a wide range of human diseases, including cancer [15], which promotes the interest in understanding their roles. A variety of functional studies in different species have shown that lncRNAs play important functions in numerous key cellular pathways, including regulation of gene expression, progression through the cell cycle, and establishment of cell identity during embryonic development [16-19]. While it remains unclear how most of these functions are carried out, it is almost certain that different lncRNAs employ radically different modes of action, some of which take place predominantly in the nucleus, and others in the cytoplasm [14]. As the biological mechanism of action of the vast majority of lncRNAs remains unknown, a lurking question is whether some of them actually encode short functional peptides, and whether some of the lncRNAs might function as mRNAs.

How can one distinguish between coding and noncoding transcripts among long RNAs? lncRNAs rarely contain highly structured regions, making tools for predicting classical structured noncoding RNAs of limited utility. Further, vertebrate lncRNAs are on average ∼1000 nt long, and so by chance they are expected to contain multiple translatable open reading frames (ORFs) with at least 50 (and sometimes even more than 100) in-frame codons enclosed within an AUG codon and a stop codon [20]. By mere chance, many of the AUG start codons in such ORFs will also be found in favorable Kozak contexts that will promote translation initiation. Therefore, according to textbook knowledge of translation, many annotated lncRNA transcripts *should* in fact be coding. The options that we have to consider for each lncRNA are therefore: (i) it indeed codes for a functional protein; (ii) it employs a mechanism that allows it to avoid productive translation; and (iii) it is translated, and translation does not result in functional peptides, and so is either tolerated by the cell as “waste” or has other roles. As we will discuss here, the relative fraction of currently known lncRNAs that can be assigned to each of the three groups is controversial, with some studies suggesting that hundreds of human lncRNAs are coding for uncharacterized proteins [21, 22], while others concluding that only very few are in fact protein-coding [23, 24]. We propose that most current evidence points to the prevalence of the third option – lncRNAs are being pervasively translated, but products of their translation are very unstable or nonfunctional.

## Computational methods for distinguishing between protein-coding and lncRNA genes

Different computational schemes can be used to assess the sequence or the evolution of an uncharacterized transcript and predict whether it is likely to encode a protein. As most features that can be used for such classification have limited discriminatory power, methods usually rely on a combination of diverse features. An overview of recently-introduced tools for distinguishing between coding and noncoding transcripts is found in **Table 1**. The following groups of features have been proposed by different studies as inputs for classification schemes distinguishing between coding and noncoding transcripts (**Figure 1**):

**Table 1.**
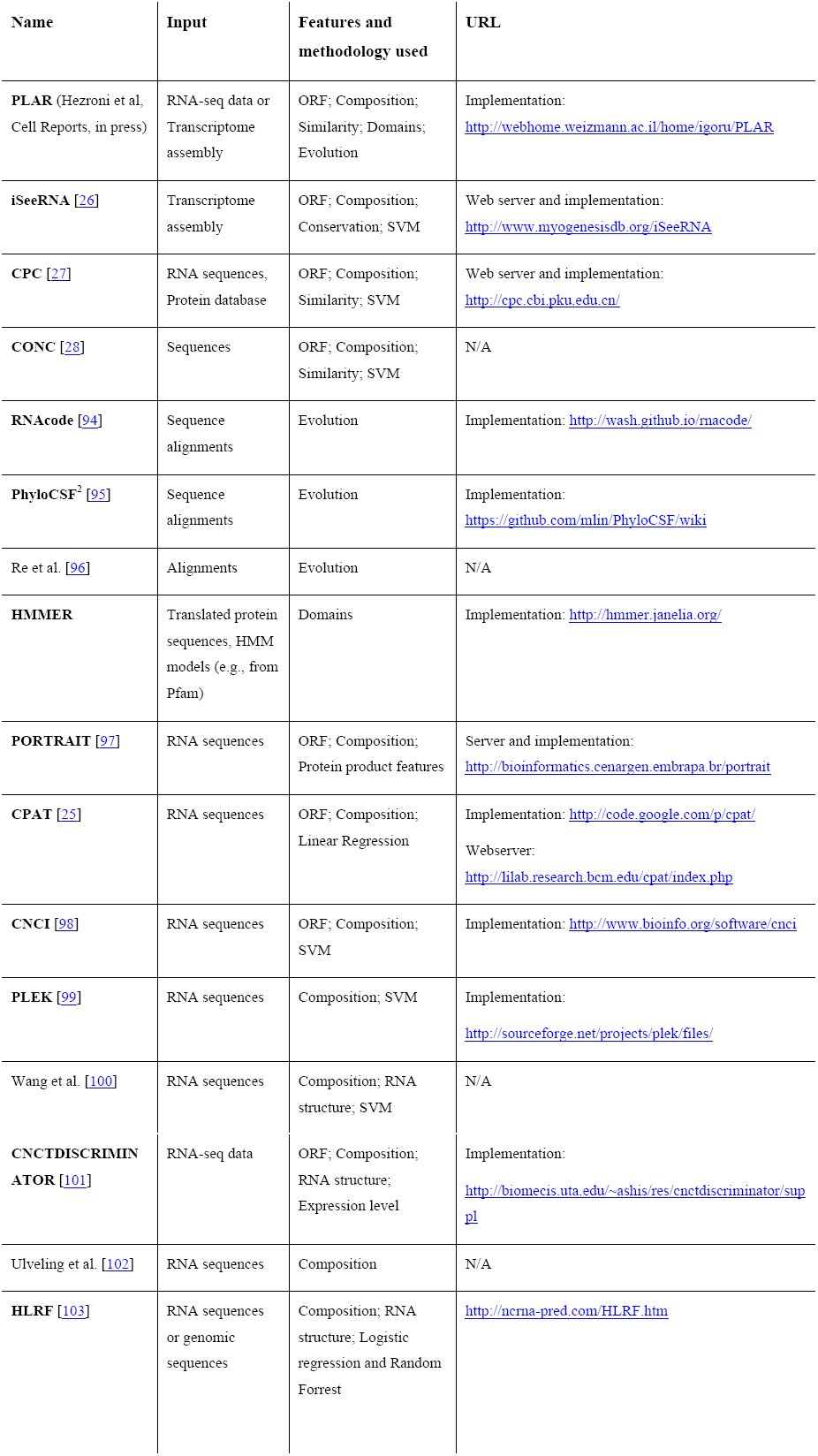
Recent methods for distinguishing between coding and noncoding transcripts. ‘Features and methodology’ refers to the types of features and means for their combination used by each method or tool: “ORF” refers to the length (absolute or relative) of ORFs in the transcript; “Composition” refers to features describing frequencies of nucleotides, amino acids or their combinations in the entire transcript or in specific ORFs; “Similarity” refers to features describing the degree of similarity between the transcripts, or its predicted ORFs, to protein sequences in known database; “Domains” refers to the potential of the transcript to encode a protein domain found in a protein domain database, such as Pfam [29]; “Evolution” refers to the substitution patterns of nucleotides in the transcript, evaluated using whole-genome alignments; “Conservation” refers to the use of general sequence conservation, e.g., computed using PhastCons [104]; “RNA structure” refers to features describing different characteristics of secondary structures predicted to form by the transcript; “SVM” refers to the use of a supporting vector machine to combine the features in a classifier framework.

**Figure 1.**
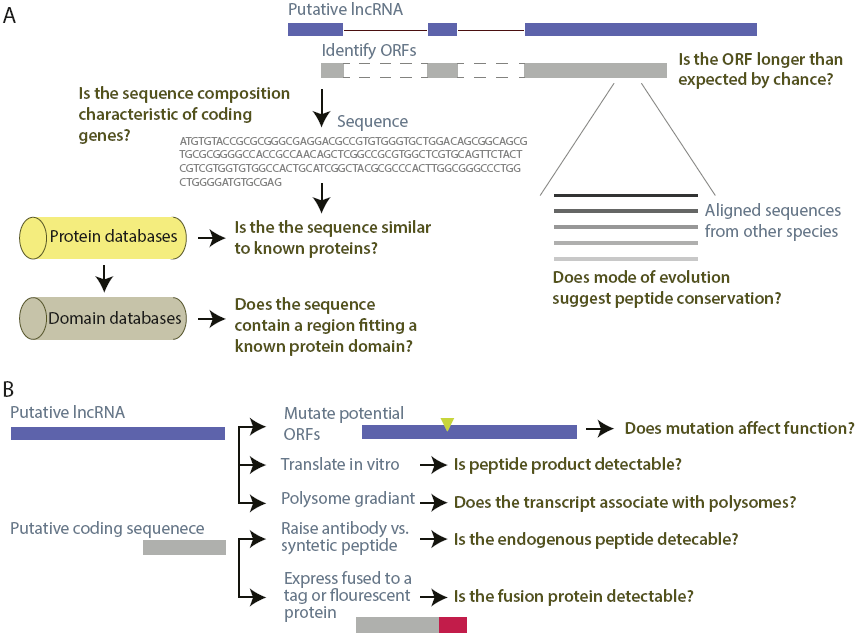
Methods for distinguishing between protein-coding and noncoding transcripts. A scheme of the common computational (A) and experimental (B) approaches for evaluating the protein-coding potential of a specific putative lncRNA transcript.

### ORF length

coding regions tend to be much longer than expected by chance [20], and so the presence of a long ORF (e.g., >300 nt long, encoding a protein with >100 amino acids) can serve as an indicator of the coding potential of a sequence. Since the likelihood of seeing a long ORF increases with transcript length, tools such as CPAT [25] and iSeeRNA [26] also examine the length of the longest ORF as a fraction of the entire transcript sequence. It should be noted that by itself, ORF length has limited predictive ability – a transcript of 2 Kb is expected to have an ORF of ∼200 nt by chance, and an ORF of 300 nt is only one standard deviation longer than expected by chance [20]. Indeed, well characterized human lncRNAs including *H19*, *Xist*, *Meg3*, *Hotair*, and *Kcnq1ot1* all have ORFs of >100 codons [20].

### Nucleotide, codon or short word frequencies

nucleotide frequencies in ORFs encoding functional proteins are dictated by non-random codon usage, and so the spectrum of nucleotide or word frequencies in entire transcripts or predicted ORFs can be used as indicators of coding potential. As shown in **Table 1,** many of the recent tools for distinguishing between coding and noncoding transcripts rely mostly on this group of features, in part because they are easier and faster to compute than features based on sequence conservation or similarity. Some tools, such as CPC [27] and CONC [28], look at triplet composition, and at the properties of the amino acids encoded by them, whereas others, such as CPAT [25], also look at frequencies of individual bases at each possible frame, and hexamer frequencies in the entire sequence.

### Substitution patterns

Protein-coding sequences evolve under selective pressures to preserve specific amino acids or amino acid types at defined positions and to maintain open reading frames. The presense of such pressures can be measured by inspection of multiple sequence alignments: by comparing substitution frequencies in different positions within a reading frame, and by testing whether insertions and deletions (indels) are depleted, and the reading frame is preferentially preserved when indels do occur. Features derived from sequence alignments are particularly powerful for detecting transcripts that encode conserved peptides, even very short ones that are difficult to recognize using other features. Naturally, such features are less useful for detecting peptides that only recently became functional, or for annotating lncRNAs in species where related genome sequences or whole genome alignments are not available.

### Presence of sequences encoding known functional domains

protein-coding genes typically contain common protein domains and the probabilistic models describing those domains have been collected in dedicated databases, such as Pfam [29] which contains Hidden Markov Model (HMM) representations of both well-characterized and putative domains [30]. Tools such as HMMER (http://hmmer.janelia.org/) can then be used to examine possible products of transcripts in all three frames and compute the likelihood that the transcript encodes a common domain, which in turn provides substantial evidence that the transcript is protein-coding.

### Similarity to known proteins

coding regions are likely to bear sequence similarities to entries in known protein databases. This feature is superficially simpler to implement than some of the other features, but it is important to carefully choose and filter the protein sequence database that the putative lncRNA is compared to, as some databases, including both GenBank and Ensembl frequently contain “hypothetical protein” sequences or models with no experimental support. It is also worth noting that true noncoding transcripts that have recently adopted sequence from a coding transcript (e.g., from a pseudogene) may contain elements that score highly as potential functional domains or as similar to other proteins even those do not reside in a coherent open reading frame.

### Other feature groups

Additional features have been proposed as suitable for classification, but those have more limited support in known *bona fide* lncRNAs. For instance, it is well appreciated that lncRNAs evolve much faster than protein coding genes, and so iSeeRNA [26] uses general sequence conservation as a criterion for distinguishing between coding and noncoding RNAs. However this feature likely introduces a bias against the rare conserved lncRNAs [9, 13]. Other tools (**Table 1**) also use the presence of structured elements (quantified as minimum free energy of the predicted fold) as features, but there is no conclusive evidence that lncRNAs are more structured than mRNAs [14, 31, 32], and in fact there are predictions that lincRNAs may even be less structured [33] and experimental data suggesting they are slightly more structured than mRNAs, but still much less structured than canonical short or intermediate size ncRNAs [34].

Most of the studies extracted some of the features described above from collections of “positive” and “negative” samples (known protein-coding sequences and some set of “confidently noncoding” RNAs) and then trained a classification algorithm, such as a support vector machine (SVM) or a logistic regression [35]. For instance, the widely used Coding Potential Calculator (CPC) [27], uses the length of the ORF in a transcript, its codon frequencies, and the BLASTP-computed similarity scores between transcript ORFs and a known protein database as inputs to an SVM [35] that outputs a binary prediction (coding/noncoding) along with a confidence score. As very few long RNAs have been rigorously shown to be *bona fide* noncoding, the most challenging aspect of such studies is the construction of the set of “negative” samples – sequences that are known with high confidence to be noncoding. The use of different such datasets in different studies makes it difficult to compare and evaluate the reported performance.

The underlying assumption behind all the criteria currently used is that short, recently evolved, yet functional proteins are relatively rare. As we will describe below, current mass spectrometry data support the notion that such proteins are rare, but the extent of their prevalence remains the pivotal question in this controversial topic.

Complementing the tools for detecting transcripts that are likely to be noncoding are methods for detection of high-confidence translated small ORFs (sORFs) (reviewed in [36]), such as of sORFinder (http://evolver.psc.riken.jp/) [37], HAltORF (http://www.roucoulab.com/haltorf/) [38] and uPEPperoni (http://upep-scmb.biosci.uq.edu.au/) [39].

## Experimental approaches for testing whether individual transcripts are translated into proteins

In pursuing the function of a specific lncRNA, even if it passes the computational filters described in the previous section, it is desirable to test experimentally whether it is indeed noncoding (**Figure 1B**). Since most transcripts have only few ORFs that are suspiciously long or contain potentially conserved amino acids, it is feasible to use several alternative methods to test if any of those are translated into detectable peptides. The best option, in our opinion, is to test whether the functionality of the transcript is preserved when the ORFs are perturbed, e.g. by introducing frameshift-inducing mutations [9, 40]. This approach is only applicable when the function of the lncRNA is known and when the transcript sequence can be manipulated (e.g., when the relevant phenotype results from transcript over-expression [40] or if a custom-made transcript sequence can be used in a rescue experiment [9]).

Each of the other available methods is associated with caveats. One can test if the transcript yields peptides when translated *in vitro* [41, 42], but a sequence may be translated *in vitro* but not *in vivo* and vice versa. Predicted peptides can also be synthetized and used to produce antibodies that can then be used to detect the putative peptide by various methods such as immunohistochemistry or western blot, though those methods may have limited sensitivity [43]. Another option is to fuse the predicted ORF with a C-terminal tag such as FLAG or eGFP that can then be used for detection using Western blotting or microscopy [43-46]. The problem here is that fusion of a peptide to such a tag can turn a very unstable peptide into a part of a stable longer protein. It is also possible to test if the transcript associates with polysomes [47, 48], but association with polysomes does not necessarily imply that the protein products are functional; indeed, transcripts considered *bona fide* lncRNAs, such as H19 and HULC, associate with polysomes [43].

### Using ribosome profiling to distinguish between coding and noncoding transcripts

A combination of the approaches listed above can lead to quite conclusive evidence about the coding potential of specific transcripts. However, the holistic question of the extent to which the currently annotated lncRNAs are “noncoding” requires globally applicable methods. Ribosome profiling [49] (Ribo-seq) allows one to take a snapshot of RNA regions that are associated with translating ribosomes. It combines classical approaches for obtaining “ribosome protected fragments” (RPFs) – footprints of actively translating ribosomes [50] – with the advances made in deep sequencing in the past decade, in order to obtain a global map of the positions within eukaryotic RNAs occupied by 80S ribosomes. Application of Ribo-seq in different systems, including mice [21], zebrafish [51], plants [52], and yeast [53], has shown that a substantial portion of expressed lncRNAs are associated with translating ribosomes. For instance, Ingolia et al. [21] found that the majority of annotated lncRNAs in mouse ES cells have regions associated with ribosomes with efficiencies closely resembling those of coding sequences and substantially higher than those observed for mRNA 3′ UTRs. The initial interpretation of this finding was that many unannotated lncRNAs are in fact translated, perhaps polycistronically, and encode multiple short peptides [21].

Subsequent studies challenged this notion. Guttman et al. [24] and Chew et al. [54] highlighted other characteristics of these data that distinguish lncRNAs from coding sequences, such as a sharp decrease in ribosome occupancy downstream to a stop codon seen in canonical coding sequences [24], but much less so in lncRNAs. This criterion, by itself or in combination with other footprint- or ORF-based features, can be used to effectively distinguish lncRNAs from canonical coding sequences, though not between lncRNAs and 5′ UTRs [24, 54].

Another important drawback raised was that some of the putative RPFs might result from RNA protection by ribonucleoprotein complexes other than translating 80S ribosomes, since some footprints were found on canonical ncRNAs that are certainly not translated such as the RNA components of RNAse P and telomerase, and since the the lengths of these footprints were different than those of RPFs from canonical coding sequences [24, 55]. The most recent reanalysis of the existing data along with new techniques introduced by Ingolia et al. [55] suggest that while such non-ribosomal footprints are indeed present in typical Ribo-seq libraries, they can be effectively removed to further enrich for reads corresponding to *bona fide* 80S footprints. One method that allows this distinction is FLOSS [55], a metric that ranks ORFs according to similarity of their ribosome footprints length distribution to the length distribution of known coding sequences. However, while FLOSS scores effectively removed footprints on mitochondrial genes and classical ncRNAs, the vast majority of the RPFs observed on lncRNAs and 5′ UTRs were still retained, reinforcing the observation that those regions were indeed actively translated. Additional experiments, including direct pulldown of ribosomal proteins added strong support to this conclusion.

The current notion is therefore that many lncRNAs undergo active translation, but that this translation resembles that observed at 5′ UTRs, in particular in that either translation termination is not very efficient or some ORFs are commonly skipped. Several recent studies combined Ribo-seq with other computational and experimental methods in order to further increase confidence in detecting translation events:

- <u>Drug treatments</u>: different drugs target translation at specific stages, and conducting Ribo-seq experiments after such treatments can yield a better understanding of the profiling, or be used for particular goals. For example, the use of heringtonine [21] or puromycine [56] uncovered unannotated translation initiation sites and the prevalence of non-AUG start codons.
- <u>Polysomal fractionation</u>: Ribosomal footprinting gives indication of what regions are translated, but not of the relative fraction of the copies of a transcript that are undergoing active translation, and polysomal fractionation can address the latter question. RNA-seq in subcellular fractions of human cells showed that lncRNAs are present in different cellular compartments – nucleus, free cytosolic, and ribosome-associated fractions, with most being enriched in the free cytosolic [57] and in the light polysomal fraction [58]. Poly-ribo-seq, i.e. ribosome profiling following polysomal fractionation, allows enrichment of transcripts that are translated by only a few ribosomes and therefore likely contain small ORFs (smORFs). Aspden et al. [59] used Poly-Ribo-seq in search of novel smORFs in flies, but generally found that smORFs in ncRNAs had profiles different than those of known protein coding genes encoding long or short ORFs. The novel smORFs had a lower translational efficiency, similar to UTRs and their products could rarely be validated with mass-spectrometry or with FLAG tag signal. Notably, the ‘dwarf’ smORFs in lncRNAs were shorter (∼20 aa) than known smORFs in flies.
- <u>Machine learning methods for filtering footprints</u>: As noted above with FLOSS, one can use metrics to classify the Ribo-seq reads into those that represent translation and those that do not. Such metrics can classify by read density similar to the density in coding regions [60], the match of the RPF to the reading frame [51], and nucleotide composition and conservation [61].

Importantly, these studies showed association of many lncRNAs with translating ribosomes and presumably the formation of translation products, but as discussed below, translation does not warrant production of a functional peptide. Indeed, for the most part, there is no current evidence indicating that more than a handful of the lncRNA-encoded peptides are stable, functional or conserved. For example, Bazzini et al. [51] used a computational approach to predict 303 coding smORFs in transcripts considered non-coding, 71% of which are under 100 aa long. However, only six novel peptide products of these smORFs, all in lncRNAs or UTRs, could be detected by mass spectrometry (see below).

### Mass spectrometry for detection of peptides from novel genes – lack of evidence or lack of power?

The next question is to what extent the peptide products of the translation events on lncRNAs accumulate in cells at detectable levels? Mass spectrometry (MS) is currently the leading proteomic platform for peptide detection and a multitude of recent studies used different flavors of MS for studying peptides expressed in human, mouse and zebrafish tissues [22, 23, 44, 51, 62-69]. *A priori*, it might be difficult to detect potential peptides originating from lncRNAs using classical MS design due to at least three issues: (i) experimental biases against detection of peptides smaller than 10 kD; (ii) paucity of potential peptides coming from unannotated transcripts in databases used for spectra search; and (iii) the inherent limited sensitivity of MS which leads to bias against detection of lowly expressed genes. Accordingly, recent studies have proposed improved MS methods, including a peptidomics approach [44, 70], combined fractional diagonal chromatography (COFRADIC) [67, 69, 71, 72], and other improvements [62, 68]; constructed potential peptide databases using RNA-seq or Ribo-seq-based transcriptome reconstructions [44, 62, 67, 73]; and combined multiple MS experiments to increase proteome coverage depth [22, 63].

Despite these advances that have increased the *a priori* chances of observing lncRNA translation products, and specific focus of recent MS-based surveys on identifying such spectra (which may have even led to some degree of ascertainment bias), the most recent attempts still found very limited evidence of peptide products that can be traced to lncRNA genes:

- As part of the ENCODE project, Banfai *et al.* [23] analyzed shotgun MS/MS data from K562 and GM12878 cell lines, and after filtering unannotated protein-coding genes found peptides mapping to only two lncRNAs, each of which was supported by just one peptide. When compared with peptide detection rates of mRNAs with expression levels matching those of lncRNAs, lncRNAs were 13- to 20-fold depleted for detected translation given their expression levels. Importantly, this study did not detect a general bias against detection of peptides coming from short ORFs [23].
- Slavoff *et al.* [44] used a peptidomics method and identified just eight peptides <50 aa derived from lncRNAs, and for those eight, the evidence that they come from mature peptides is limited [74].
- Kim *et al.* [63] identified nine annotated non-coding peptide-producing transcripts in a very deep proteomic survey (16 million MS/MS spectra). The same group reported peptides from 34 lncRNAs in zebrafish [66], where a higher number of mis-annotated lncRNAs is expected due to less mature genome sequence and annotation.
- Bazzini et al. [51] used a dedicated MS approach for identifying protein-products of zebrafish small ORFs predicted based on conservation and ribosome profiling (see above), but found peptides from products of just six new lncRNA ORFs.
- Prabakaran *et al.* [73] reported 250 novel peptides coming from unannotated regions, but only 25 of those mapped to regions outside the boundaries of protein-coding genes, and only three of those had support from Ribo-seq data. When we mapped the 250 peptides from this study against a collection of lncRNAs defined by PLAR (**Table 1**) and passing stringent filters, we found only two matches, and both mapped to probable pseudogenes.
- Another study specifically designed to detect peptides coming from unannotated genes or lncRNAs did not detect any evidence of translation from lncRNAs [64].
- Finally, several other recent studies did not report any peptides derived from lncRNAs, despite the use of a Ribo-seq-based peptide database [69, 72].

Notably, the ability to detect translation products derived from translation events in 5′ UTRs was also very limited, and few of the ORFs that appeared confidently translated based on ribosome footprinting were detected by MS, suggesting similarities between translation outcomes in lncRNAs and 5′ UTRs [51, 67].

The studies described above appear to contrast a recent study by Wilhelm *et al.* [22] that reported 430 peptides from 404 lncRNAs detected by MS. This unexpectedly high number suggested that many lncRNAs may indeed be translated and produce detectable peptides, that were somehow missed by others. However, our analysis of the data suggests that these large numbers result from promiscuous search parameters when matching spectra to potential transcripts. In fact, when we used BLASTN to map the 430 peptides from Wilhelm *et al.* [22] to lncRNA exons in our recent collection of >10,000 human lncRNA genes passing stringent filters (Hezroni et al., *Cell Reports*, in press), we found only five hits with BLAST E-value < 10^-3^, and all of those mapped to pseudogenic regions. Our analysis is concordant with a recent re-analysis of these data by Valencia, Tress, and colleagues [75] that concluded that the peptide identifications reported by Wilhelm *et al.* [22] likely contain many false positives due to inclusion of low-quality spectra and relaxed peptide detection thresholds.

We conclude that evidence from about a dozen recent studies suggests that translation products originating from ORFs in lncRNAs, even those that appear translated in Ribo-seq data, are essentially invisible in MS data. While it is possible this is due to limitations of MS in analyzing peptides shorter than 50 aa [51], we propose that a more likely explanation is that such peptides are very unstable and therefore do not accumulate to consequential levels in mammalian cells.

## Discussion

### Best practices for annotating a transcriptome

As described above, the toolbox for distinguishing between coding and noncoding transcripts is substantial and rapidly growing, and many of the tools are available both as web servers and as stand-alone tools (listed in **Table 1**). Some of these tools (typically those that rely only on sequence composition features) are relatively simple to use, while others require dedicated input processing. In our experience each of the tools has limitations when used in isolation, and so better accuracy is achieved by combining multiple methods, as we have recently done as part of our Pipeline for lncRNA annotation from RNA-seq data (PLAR, Hezroni et al., Cell Reports, in press). It also remains very important to manually inspect transcripts of particular interest even if they are predicted as coding, and test whether the evidence of protein-coding potential is not a result of an artifact, and indeed converges on a specific ORF that is likely to be coding. For example, sequences that are not repeat-masked will sometimes contain regions resembling protein domains, and regions that are very highly conserved will sometimes be called coding simply because their alignments are not informative enough for tools inspecting substitution patterns.

It should be also be noted that, in our experience, when studying a well-annotated genome, such as human or mouse, only few RNA-seq reconstructed transcripts that are classified as coding by the above-mentioned but that do not overlap known genes actually correspond to *bona fide* unannotated proteins. Rather, the majority of such transcripts are usually parts of pseudogenes, which are sometimes difficult to identify [76]. It is also notable that some recently described lncRNAs with specific functions fail one or more of the commonly used filters. For example, TINCR [77] contains a region predicted by RNAcode to be coding and the region covering its 456 nt ORF is predicted to encode a peptide that matches an uncharacterized Pfam-B domain. Another lncRNA, lnc-DC, is a human ortholog of the mouse protein-coding gene 110000G20Rik which encodes for a Wdnm1-like protein (Wfdc21) that is highly conserved in mammals [78], and therefore would not pass protein-similarity filters.

### Functionality of pervasive translation of lncRNAs?

The question of the functionality of translation in lncRNAs can be reduced to the question of the extent to which peptides <50 aa, after their release from the ribosome, can remain stable and functional in vertebrate cells. Inspection of proteins with currently known functions that are annotated in Ensembl reveals only 16 that are <50 aa. Some of these are clearly independently translated proteins, such as Ost4, a 37aa, 3.4 kD membrane protein conserved all the way to yeast. Such short proteins can therefore be stable and functional in vertebrates, but their relative scarcity indicates that they may be the exception rather than the rule. We suggest that the vast majority of other short-peptide translation events formed as part of “pervasive translation” [55, 79], which appear very common based on Ribo-seq data, are largely non-functional products that are tolerated in cells because they are usually rapidly degraded by abundant cytosolic endopeptidates, aminopeptidases, and other machineries active in cells [80] and therefore have limited impact. Systematic evidence for stability of short (<50 aa) peptides in cells would be invaluable for addressing this question, but no such resources are presently available, and existing anecdotal evidence suggests that such peptides are generally very unstable [80-83].

When the cost of production of such peptides is considered, it is important to keep in mind that the vast majority of such translation events occur in 5′ UTRs of coding genes rather than in lncRNAs. Using Ribo-Seq data from U2OS cells [84] we find that a similar fraction of all mapped RNA-seq reads (and by proxy nucleotides in polyadenylated transcripts) are found in 5′ UTRs and lncRNAs, but 5′ UTRs account for 10-times more Ribo-seq reads (and by proxy for 10-times more ribosomes) (**Figure 3**). When comparing to coding regions and using the Ribo-seq reads as a proxy for the number of ribosomes actively translating lncRNAs and mRNAs, there are over 1,000-times more ribosomes associated with mRNAs than with lncRNAs. Similar results are obtained in other cell types (I.U., unpublished data). The total fitness cost of all the translation events from all lncRNAs is thus small when compared to the cost of mRNA translation.

**Figure 2.**
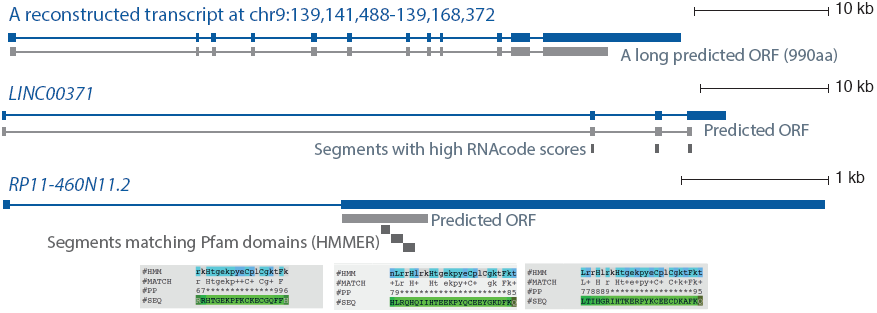
Examples of putative human lncRNA transcripts that do not pass one or more of the commonly used filters for detecting protein-coding genes. The first transcript is reconstructed from human RNA-seq data by cufflinks (Hezroni et al, Cell Reports, in press). And contains a long ORF. The second is presently annotated as a lincRNA but overlaps several domains predicted to be coding by RNAcode [94]. The third encodes an ORF that is predicted by HMMer to encode zinc-finger domains annotated in the Pfam database.

**Figure 3.**
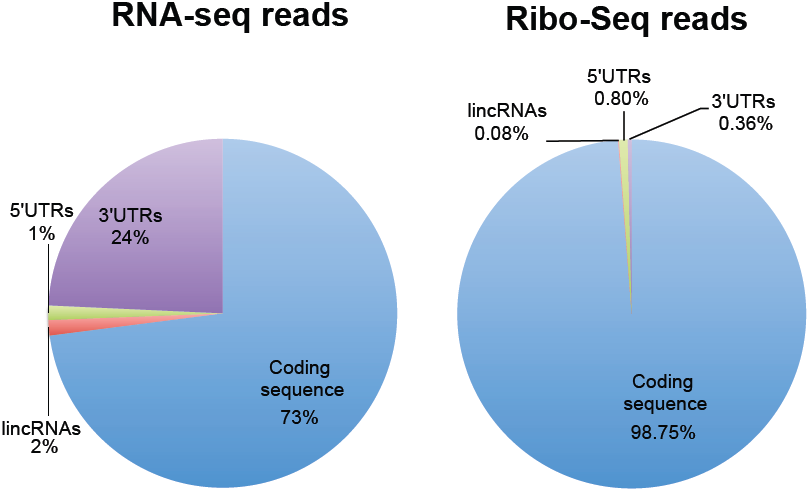
Distribution of RNA-seq and Ribo-seq reads. RNA-seq and Ribo-seq data from human osteosarcoma U2OS cells [84] were mapped to the human transcriptome, and only reads mapping to mRNAs or lncRNAs were considered. Plotted is the fraction of reads mapping to the indicated features.

A different and interesting way to view pervasive translation of lncRNAs is that these genes can serve as a platform for de novo protein evolution [60, 85]. Characteristics of a majority of lncRNAs, in particular, their relative novelty and lineage specificity, suggest that these lncRNAs could be strong candidates as precursors for new proteins. Furthermore, new proteins are expected to be very short and under weak evolutionary constraints, fitting the translated smORFs identified in lncRNAs.

### Regulatory roles of translation in noncoding transcripts

The fate of the translation products arising from lncRNAs is part of a bigger question of translation of “untranslated regions”, and among those most prominently of 5′ UTRs. Since mRNAs as a group are ∼100-fold more abundant than lncRNAs in cells, and most 5′ UTRs have at least one uORF that undergoes canonical translation, 5′ UTRs are much more abundant templates for translation of small ORFs than lncRNAs, yet the products of this translation are also almost entirely absent from MS data (as detailed above) and the ORFs that are being translated are only very rarely conserved in sequence. Very few ORFs in 5′ UTRs are conserved between human and mouse, and even in those that are conserved, selection on the encoded peptide sequences is mostly weak [86].

It is not known how much of the translation observed in uORFs is functional and how much is tolerated noise of the translational machinery, but some of this translation functions in regulating the translation of the main (almost always longest) ORF in the mRNA [87, 88]. We can envision parallel “regulatory” roles for translation of ORFs in lncRNAs. Specific possible regulatory roles include regulation of stability of the lncRNA (e.g., by modulating its degradation by nonsense mediated decay [89, 90] or other pathways), regulation of its localization in the cell, or protection of parts of the lncRNA sequence from scanning ribosomes that are potent helicases. Translation-facilitated degradation of long noncoding RNAs can serve as an elegant mechanism for regulating the accumulation of snoRNA precursors, which is further modulated during stress [91-93] – and so translation can certainly induce a regulatory mechanism in long noncoding RNAs. We argue that such mechanisms are probably much more prevalent than currently appreciated and should serve as a fruitful direction of further research into the biological consequences of pervasive translation of lncRNA genes.

## Acknowledgements

We thank Rory Johnson, Yoav Lubelsky, Lisha Qiu Jin Lim, Noa Gil, Hadas Hezroni and Neta Degani for useful discussions and comments on the manuscript. I.U. is incumbent of the Robert Edward and Roselyn Rich Manson Career Development Chair and recipient of an Alon Fellowship. Work in the Ulitsky lab is supported by grants to I.U. from the Israeli Science Foundation (1242/14 and 1984/14), the Minerva Foundation, the Fritz-Thyssen Foundation and the Rising Tide foundation.

## Footnotes

1 Uses CPC, HMMER, RNAcode and combines their results.

2 Parameters currently available only for whole-genome alignments for mammals, flies, mosquitos and yeast.

